# Mix & Match: Phenotypic coexistence as a key facilitator of solid tumour invasion

**DOI:** 10.1101/750810

**Authors:** Maximilian A. R. Strobl, Andrew L. Krause, Mehdi Damaghi, Robert Gillies, Alexander R. A. Anderson, Philip K. Maini

## Abstract

Invasion of healthy tissue is a defining feature of malignant tumours. Traditionally, invasion is thought to be driven by cells that have acquired all the necessary traits to overcome the range of biological and physical defences employed by the body. However, in light of the ever-increasing evidence for geno- and phenotypic intra-tumour heterogeneity an alternative hypothesis presents itself: Could invasion be driven by a collection of cells with distinct traits that together facilitate the invasion process? In this paper, we use a mathematical model to assess the feasibility of this hypothesis in the context of acid-mediated invasion. We assume tumour expansion is obstructed by stroma which inhibits growth, and extra-cellular matrix (ECM) which blocks cancer cell movement. Further, we assume that there are two types of cancer cells: i) a glycolytic phenotype which produces acid that kills stromal cells, and ii) a matrix-degrading phenotype that locally remodels the ECM. We extend the Gatenby-Gawlinski reaction-diffusion model to derive a system of five coupled reaction-diffusion equations to describe the resulting invasion process. We characterise the spatially homogeneous steady states and carry out a simulation study in one spatial dimension to determine how the tumour develops as we vary the strength of competition between the two phenotypes. We find that overall tumour growth is most extensive when both cell types can stably coexist, since this allows the cells to locally mix and benefit most from the combination of traits. In contrast, when inter-species competition exceeds intra-species competition the populations spatially separate and invasion arrests either: i) rapidly (matrix-degraders dominate), or ii) slowly (acid-producers dominate). Overall, our work demonstrates that the spatial and ecological relationship between a heterogeneous population of tumour cells is a key factor in determining their ability to cooperate. Specifically, we predict that tumours in which different phenotypes coexist stably are more invasive than tumours in which phenotypes are spatially separated.

## 1 Introduction

Tissue invasion is a hallmark of cancer [31]. If a tumour is detected before it has started to spread into the surrounding tissue then the tumour is termed *benign* and the chances of survival are high. If the tumour has started to spread, breaching the basement membrane, survival rates are significantly decreased and the tumour is termed *malignant* (“badly born”). In 90% of patients the cause of death is not the primary tumour, but the disruption of normal body function caused by metastatic disease [46] - for which invasion is the first critical step.

Due to the profound damage caused by the uncontrolled spread of cells, a great number of mechanisms have evolved to ensure that cells - even those that might have started to escape homeostatic control - remain localised. One important barrier, for example, is the extra-cellular matrix (ECM), a dense mixture of proteins encapsulating the cells in healthy tissue [47, 53]. The proteins in the ECM form a strong scaffolding which physically anchors tissue cells in place and activates intra-cellular signalling pathways which suppress cell movement and regulate proliferation [47, 53, 37, 53, 9]. A further important barrier to local expansion of the tumour is the inhibitory environment created by the healthy tissue (stroma) surrounding the tumour. For example, an analysis of 432 different cancer-fibroblast co-cultures found that 41% of the investigated pairings led to reduced cancer growth [50].

Research over the past decades has elucidated in great detail the molecular mechanisms used by cancer cells to overcome these barriers. In order to remodel or degrade the ECM, tumour cells use matrix degrading enzymes (MDEs) such as matrix metalloproteinases [47, 16, 31]. Similarly, in order to overcome the growth inhibition from the surrounding stroma, tumour cells can coerce healthy cells into tumour promoting phenotypes (e.g. tumour associated fibroblasts), or eradicate them. In a series of papers, Gatenby and co-workers have proposed that an important contribution to this transformation is the acidification of the tissue environment by the tumour, a theory known as the “acid-mediated invasion hypothesis” [24, 26, 25, 30]. Many invasive cancers are characterised by their use of glycolysis for energy generation even in conditions under which more efficient aerobic respiration would be feasible, a paradox known as the “Warburg effect” [51, 30]. Gatenby and co-workers argue that the acidification due to upregulated glycolysis, which ranges over 0.5-1 pH units [54, 32], results in death of normal cells, thereby allowing tumour cells to expand [24, 26, 25, 30]. This hypothesis is supported, for example, by experiments showing that low pH leads to increased rates of apoptosis [49] or that administration of a neutralising buffer can reduce tumour expansion in mice [26].

The advances in our molecular understanding of invasion have been accompanied by a significant body of theoretical work that has aimed to integrate the insights from different spatial and temporal scales to identify clinical implications and to guide future experiments (see [4] for an excellent review). Gatenby and Gawlinski developed a mathematical model based on reaction-diffusion equations to investigate the feasibility and implications of the acid-mediated invasion hypothesis [24]. In their three-compartment model, the authors represent tissue as a mixture of healthy stromal cells, cancer cells, and acid released by the tumour cells [24]. They identify different modes of invasion depending on the system parameters, and predict that particularly aggressive invasion gives rise to a gap between the advancing tumour and retreating tissue front [24]. Subsequent work has more formally analysed this model and suggested new experiments that could be used to test the underlying assumptions [22, 36].

In addition to the role of acid, the dynamics of ECM remodelling and degradation has also been studied. Considering the ECM as a purely physical barrier, Martin and co-workers [35] used an extension of the Gatenby-Gawlinski model to demonstrate that if a collaboration between the tumour cells and the stroma is required to degrade the matrix, then highly acidic tumours may be encapsulated and unable to invade. Other studies instead considered the stimulatory effects that certain by-products of matrix degradation have on activation and direction of tumour cell movement. Anderson *et al* [2] showed in a partial differential equation (PDE) model that such an ECM gradient driven migration (haptotaxis) can influence the shape of the growing tumour. In a series of papers, the group led by Mark Chaplain have further characterised the importance of cell-cell adhesion in tumour invasion [11, 28, 18] and identified the plasminogen urokinase activation system as a key driver of invasion [14, 43, 1].

While we have an increasing understanding of *how* tumour cells invade, an important open question remains as to *when in oncogenesis* invasion emerges. Traditionally, invasion is thought to be carried out by a subset of cancer cells that have acquired all the necessary traits to overcome the host’s various defence mechanisms. However, over the past decade it has become clear that tumours are a heterogeneous mixture of cells that differ in their genetic make-up and phenotypic behaviour [38, 29, 7]. As part of a recent study, currently in preparation for publication [17], we observed significant heterogeneity in the distribution of matrix remodelling activity and acid adaptation amongst cancer cells in human breast cancer ducts (Figure 1). Even along the invasive front the overlap of the regions of acid production and matrix remodelling is not complete (Figure 1B). While further experimental work will be required to ratify these observations, they led us to ask the question: Instead of being driven by group of “super-cells”, could cancer invasion rather be an emergent property of cooperating specialist cells?

**Fig. 1.**
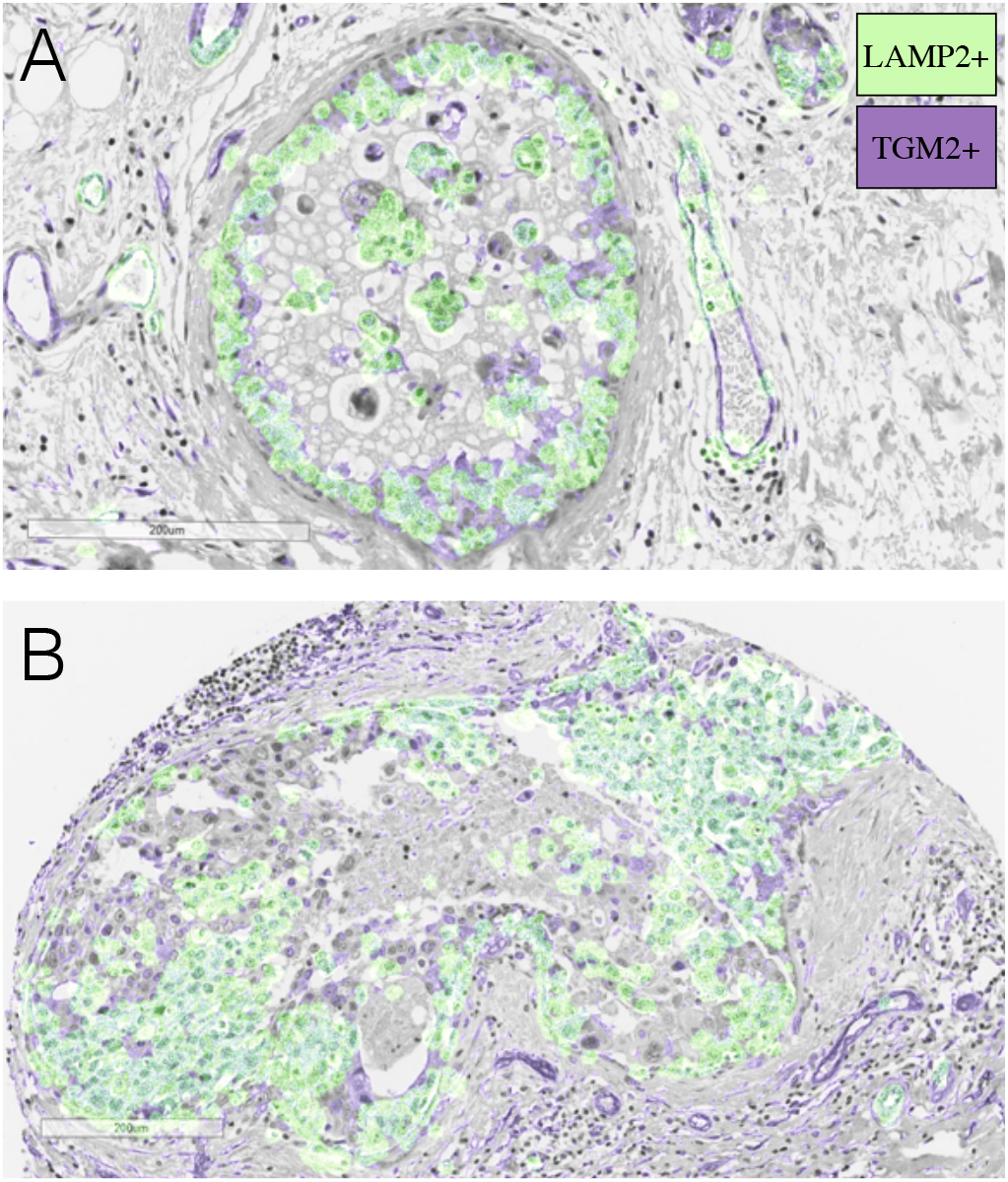
Areas of acid production and matrix re-modelling in human breast cancer ducts. Acid production was defined by expression of the acid adaptation marker LAMP2. Matrix re-modelling was defined by expression of TGM2. For visualisation purposes masks were extracted and overlaid on a haemotoxylin and eosin stain of the same tissue (see Section A1 for details). A) Example of a ductal carcinoma *in situ* that has not yet invaded the surrounding tissue. B) Example of an invasive cancer that has breached the duct. We observe that not all cells are expressing LAMP2 or TGM2. Could there be cooperation between cells with different traits?

There is mounting evidence for cooperation among tumour cells [6, 5]. A Wnt1-driven mouse model of breast cancer, for example, has been shown to be composed of two cell types: one expressing Wnt1, and the other expressing the associated Lrp5 receptor [34, 15]. Interaction with the other cell type allows each population to grow faster and drives tumour growth [34, 15]. Alternatively, production of diffusible growth factors can allow for cross-feeding amongst tumour cells, where a cell produces one type of growth factor and receives the others from its neighbours [6, 5]. Given cells have been shown to support each others’ growth, it seems plausible that they may also cooperate to overcome the body’s defences during tissue invasion.

The aim of this paper is to use a mathematical model to investigate the feasibility and the implications of this hypothesis in the context of acid-mediated invasion. We will extend the Gatenby-Gawlinski model so that it includes obstruction both from the stroma and the ECM. Specifically, we will assume that stromal cells suppress growth, whilst the ECM blocks cell movement. Unlike previous work [43, 35] we will assume that no single tumour cell can remove both obstructions. Instead, we will assume that there are two cancer phenotypes: i) an acid-producing phenotype which removes stroma, and ii) an ECM-degrading phenotype. We will assume that these distinct phenotypes cooperate to remove obstructions, but must also compete with one another for resources. Through linear stability analysis and numerical simulations of the resulting system of five differential equations we will study under which circumstances a mixture of the two populations (as defined by the relative inter-species competition) develops into an invasive cancer.

## 2 The Mathematical Model

Our model builds on the Gatenby-Gawlinski model [24], and consists of five components: stroma (*S*(**x**, *t*)), ECM (*M* (**x**, *t*)), a population of acid producing tumour cells (*T_A_*(**x**, *t*)), lactic acid (*L*(**x**, *t*)), and a population of matrix degrading tumour cells (*T*_*M*_ (**x**, *t*)), where **x** denotes space and *t* represents time (Figure 2). Following [24] we assume that densities are large enough to be describable by continuous functions, and model the spatio-temporal evolution of the system using a combination of spatially-distributed ordinary differential equations (ODEs) and PDEs.

**Fig. 2.**
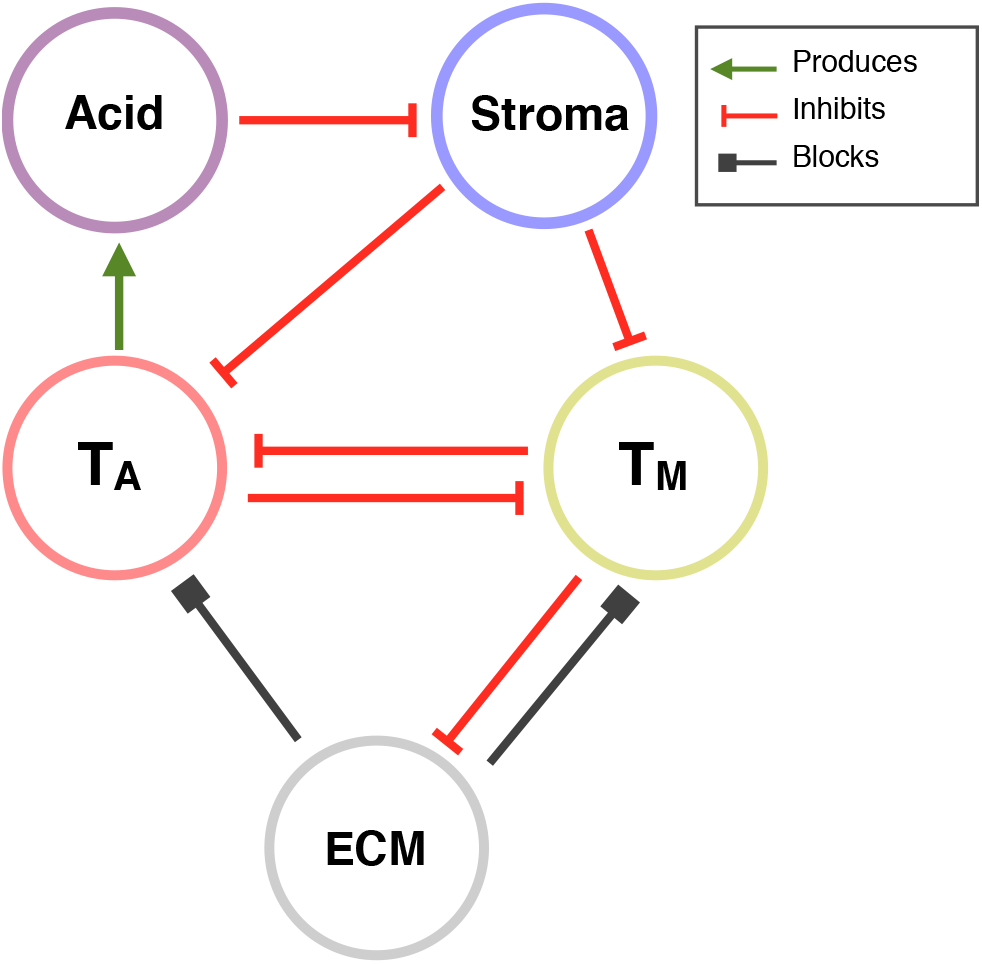
Interaction diagram of our model. Stroma inhibits tumour cell proliferation but is killed by acid secreted by the acid producing tumour cells, *T*_*A*_. ECM blocks movement of the tumour cells, but can be removed by the matrix degrading tumour cells, *T*_*M*_. The two types of tumour cells compete for resources thereby inhibiting each other’s growth.

### 2.1 Healthy Tissue Components

We consider two components of healthy tissue: stroma and ECM. The model for the stroma, denoted as *S*, is taken from [25] and assumes that:

– Stromal cells grow logistically at a rate *r*_*S*_ and carrying capacity *K*_*S*_ in the absence of tumour, reflecting homeostasis.
– Stromal cells are anchored in place and their motility can be neglected.
– Stromal cells are killed by the lactic acid produced by the tumour cells at a rate proportional to its concentration, *L*(**x**, *t*), with a constant of proportionality *d*_*S*_.

This yields the following governing equation for *S*(**x**, *t*):

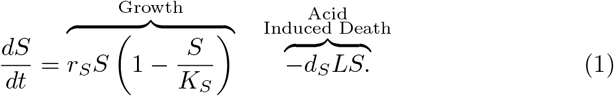

In modelling the ECM dynamics we assume:

– There is a net loss of ECM over the time-frame of interest. Since we are interested in studying the dynamics of invasion, we will assume that the break down of matrix by the tumour overcomes any regeneration, as has been done in other work previously (e.g. [42, 52, 35]).
– ECM degradation or remodelling is a localised process. This is based on the fact that MDEs are either directly located on the cell membrane or are so large that their diffusion coefficients are very small [53]. We model this with a linear mass-action model such that ECM is degraded at a rate proportional to the density of matrix degrading tumour cells, *T*_*M*_(**x**, *t*), with *d*_*M*_ the rate of degradation.
– Because of its fibrous nature, diffusion of ECM can be neglected.

Thus, we model the *M*(**x**, *t*) dynamics as:

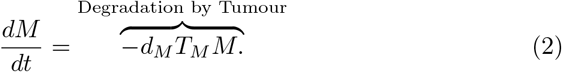

Similar models have previously been used in [2, 35]. We remark that both (1)–(2) are ordinary differential equations, that are distinct for every spatial point. Furthermore, the equations between neighbouring spatial points are not directly coupled. Instead coupling occurs via one of the other variables (e.g. *T*_*A*_).

### 2.2 The Tumour Environment

We consider two phenotypically distinct tumour populations: i) glycolytic, acid producing cells (*T*_*A*_(**x**, *t*)), which release lactic acid killing stromal cells, and ii) matrix degrading tumour cells (*T*_*M*_(**x**, *t*)), which degrade the ECM. We assume that:

– In the absence of other cells, each tumour population grows logistically at rates 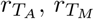 and carrying capacities 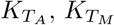 respectively.
– The tumour cells compete with each other and with the stroma for resources and space. We assume competition follows a generalized Lotka-Volterra functional response [39], characterised by competition parameters *c*_*i*,*j*_. These describe the inter-species competition that species *j* experiences from species *i* relative to the intra-species competition *i* exerts on itself.
– The tumour cells are motile, but their movement is restricted by the physical obstruction of the ECM. Following Martin et al [35], we model obstruction by the ECM as a linear reduction in the flux of cells. We denote by 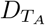 and 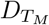 the diffusive fluxes in the absence of ECM, and by *K*_*M*_ the density of ECM such that tumour motility ceases.
– Tumour cells are resilient to the acid. Histology shows adaptation of tumour cells to acidic environments [24, 26] and theoretical work supports that acid resistance is acquired early in oncogenesis [44].

We note that our assumptions about the interactions between the tumour cells and their environment differ to those made by Gatenby and Gawlinski [24] and Martin and co-workers [35], on whose work our study is built. Specifically, Gatenby and Gawlinski choose to neglect competition between tumour and stroma, and Martin et al include the stroma as an additional physical obstruction to movement. Our choice of assumptions is motivated by the aim to make the two barriers act orthogonally, so to compare their effects. As it appears easier for cells to squeeze through the stroma in a migrating manner than the ECM we choose this particular order. Determining which assumptions are more physiologically realistic will require further study, but we anticipate that the results presented below will motivate such investigations.

In summary, we propose the following model equations for *T*_*A*_(**x**, *t*) and *T*_*M*_(**x**, *t*):

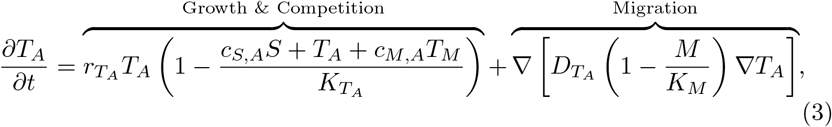

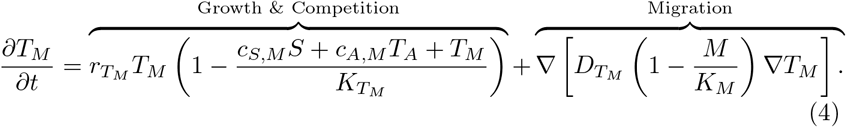

The final governing equation is that for the acid. We adopt the model by [24], and assume that:

– The acid is produced by the glycolytic phenotype, *T*_*A*_, at constant rate *r*_*L*_.
– Acid is removed from the tissue by blood vessels and natural buffering agents at constant rate *d*_*L*_.
– Because of its small molecular size, acid can diffuse unobstructedly.

This yields the following PDE for *L*(**x**, *t*):

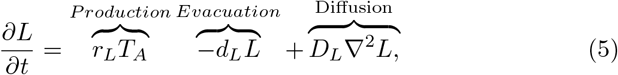

where *r*_*L*_ is the acid production rate, *d*_*L*_ the degradation rate, and *D*_*L*_ the diffusion constant.

### 2.3 Further Simplifying Assumptions

The aim of this paper is to investigate competition and cooperation between tumour cells based on distinct phenotypic properties. Thus, we will make the simplifying assumption that the two tumour populations are biologically identical, except in their abilities to degrade matrix and produce acid. We will assume identical growth rates, 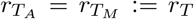, identical carrying capacities (corresponding to intra-species competition), 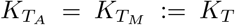 and identical motility, 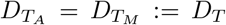. Moreover, we will assume that inhibition received from the stroma is equal for both phenotypes, so that *c*_*S,A*_ = *c*_*S,M*_ := *c*_*S*_. Finally, we will also adopt the assumption made by Gatenby and Gawlinski [24] that stroma and tumour cells have the same carrying capacities *K*_*T*_ = *K*_*S*_ := *K*.

We will study the model on a 1-d slice of tissue *Θ* = [0, *ℓ*], where *x* = 0 is the position of the initial core of the tumour and *ℓ* is the length of the slice.

We assume that the tumour has initially infiltrated a distance *σ* < *ℓ* which we model by the following initial data:

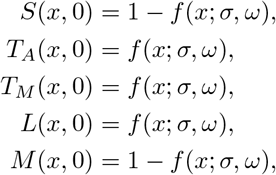

where *f* (*σ, ω*) is a mollified step function and *ω*, a fixed positive constant, describes the sharpness of the initial boundary between the tumour and the healthy tissue. Specifically:

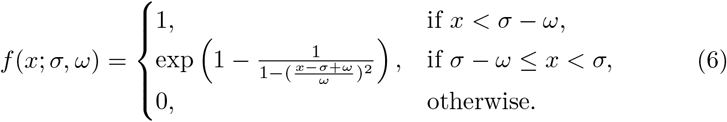

To facilitate numerical simulation we follow previous work (e.g. [24, 35]) in assuming that there are hard boundaries at *x* = 0 and *x* = *ℓ*, which allows us to close the system with zero-flux boundary conditions (at *x* = 0, *ℓ*). However, as the choice of the domain *Θ* is motivated more by numerical convenience than biological reality, we will only simulate this system for as long as the tumour is far away from the right boundary, to avoid introducing any boundary condition artefacts.

### 2.4 Non-Dimensionalisation

We introduce the following scalings, adopted from [24] and motivated by the natural scales present in the system:

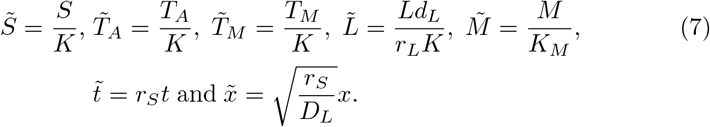

Based on the parameters used in this study (see also Section 2.5), this corresponds to a time scale of 11.57 days and a spatial scale of 2.24 cm. Following previous work [24, 35, 36] we choose *C* such that 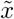 ranges from 0 to 1 for convenience. Preliminary simulations showed that this allows us to simulate for a time frame of > 600 days for most parameter combinations before the tumour starts interfering with the right boundary, which is a clinically realistic time scale (equivalent to 1 cm of tumour growth).

Dropping the ∼ for notational convenience, the re-scaled model reads:

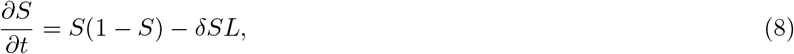

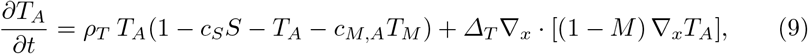

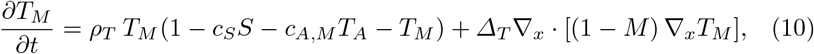

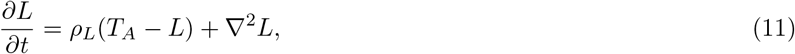

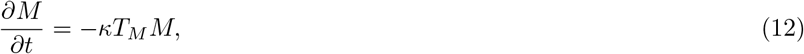

where the dimensionless parameters are given by:

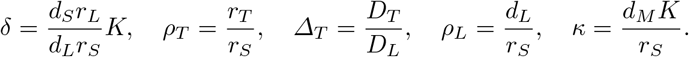

### 2.5 Parameters

As far as possible we take parameters obtained from the literature. A summary of all the parameters is shown in Table 1. The value for *κ* was adapted from [2] where it represents the maximum rate at which the MDEs can degrade the ECM. We carry out parameter sweeps in the competition parameters, as these are difficult to estimate from existing data. As we will see, the choices of ranges for competition parameters encapsulates all of the behaviours we would expect from such a model, and simulations outside these ranges can be inferred from our results.

**Table 1.** Parameters used in the numerical simulation of the model.

| Parameter | Value | Reference |
| --- | --- | --- |
| $\delta$ | 12.5 | [24] |
| $\rho_T$ | 1 | [24] |
| $\Delta_T$ | $4 \times 10^{-5}$ | [24] |
| $\rho_L$ | 70 | [24] |
| $\kappa$ | 10 | [2] |
| $c_S$ | 1.5 | Estimated |
| $c_{M,A}$ | 0-2 | Estimated |
| $c_{A,M}$ | 0-2 | Estimated |

## 3 Steady State Analysis

During invasion, tumour cells arrive in healthy tissue and establish a self-sustaining population. In principle this corresponds to a travelling wave solution (TWS) to Equations (8) – (12) which connects two spatially homogeneous steady states: the state (*S, T*_*A*_, *T*_*M*_, *L*, *M*) = (1, 0, 0, 0, *M*\*), where *M*\* is an arbitrary level of the matrix density (henceforth referred to as SS0), representing healthy tissue and another spatially homogeneous state (*S, T*_*A*_, *T*_*M*_, *L*, *M*) which describes the composition of the invaded tissue. Neglecting the trivial steady state where all cell populations are extinct, the system admits six further steady states:

– <u>SS 1:</u> (*S, T*_*A*_, *T*_*M*_, *L*, *M*) = (0, 1, 0, 1, *M*\*), which represents a tumour composed only of acid producing cells, *T*_*A*_.
– <u>SS 2:</u> (*S, T*_*A*_, *T*_*M*_, *L*, *M*) = (0, 0, 1, 0, 0), which describes a tumour composed only of matrix degrading cells, *T*_*M*_.
– <u>SS 3:</u> 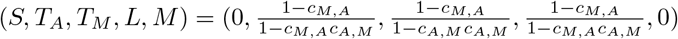, which describes cancerous tissue in which *T*_*A*_ and *T*_*M*_ coexist. Both the stroma and the ECM have been eradicated.
– <u>SS 4:</u>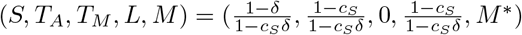, which models a tumour composed of a mixture of acid producing cells, stroma and ECM.
– <u>SS 5:</u> (*S, T*_*A*_, *T*_*M*_, *L*, *M*) = (1, 0, 1 − *c*_*S*_, 0, 0), which is representative of a tumour consisting of a mixture of matrix degrading cells and stroma. As we assume that *c*_*S*_ > 1, this state is never feasible and so not relevant to this study.
– <u>SS 6:</u> If (*c*_*M,A*_, *c*_*A,M*_) = (1, 1), (*S*, *T*_*A*_, *T*_*M*_, *L*, *M*) = (1 − *δT*_*A*_, *T*_*A*_, 1 − *c*_*S*_ − (1 − *c*_*S*_*δ*)*T*_*A*_, *T*_*A*_, 0), where *T*_*A*_ ∈ (0, 1). Otherwise, 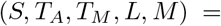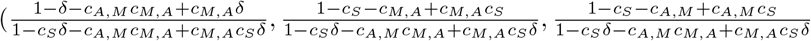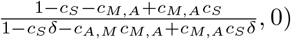. As such, SS6 represents acidic cancerous tissue in which all three cell populations coexist. The matrix has been degraded.

A linear stability analysis shows that all steady states involving a non-zero density of ECM (SS1 and SS4) have a zero eigenvalue (for details see Appendix A2). This corresponds to a perturbation in the ECM density and reflects the fact that, in the absence of *T*_*M*_, the ECM density will remain constant and all values for *M* are admissible as steady states. Furthermore, we find that SS2, SS4 and SS6 always have at least one eigenvalue with positive real part for the range of parameters considered (Figure A1). This implies that these states can not be part of a TWS representing an invading tumour. In contrast, SS1 is linearly stable if *c*_*A,M*_ > 1, whereas SS3 is stable if *c*_*A,M*_, *c*_*M,A*_ < 1 (assuming *δ* > 1; Appendix A2). We conclude that there are four possible scenarios for invasion:

1. <u>Stable Coexistence:</u> If *c*_*A,M*_ < 1 and *c*_*M,A*_ < 1, then both tumour populations stably coexist inside the tumour (SS3), resulting in an invading tumour corresponding to a TWS connecting SS3 and SS0.
2. <u>Competitive Exclusion of *T*_*A*_:</u> If *c*_*A,M*_ < 1 and *c*_*M,A*_ > 1, then *T*_*M*_ drives *T*_*A*_ to extinction inside the tumour. Where stroma is present the healthy tissue is restored (SS0), where it is absent the system settles into a monoculture of *T*_*M*_ (SS2). The tumour becomes encapsulated and invasion halts.
3. <u>Competitive Exclusion of *T*_*M*_:</u> If *c*_*A,M*_ > 1 and *c*_*M,A*_ < 1, then *T*_*A*_ drives *T*_*M*_ to extinction inside the tumour (SS1). While this might nevertheless give rise to a TWS we conjecture that the associated speed of invasion is zero due to the obstruction from the matrix. We provide numerical evidence for this in Figure 3.
4. <u>Bi-Stability:</u> If *c*_*A,M*_ > 1 and *c*_*M,A*_ > 1, then the system is bi-stable and the outcome of invasion is dependent on the initial conditions. We explore this case numerically in Section 4.2.3.

**Fig. 3.**
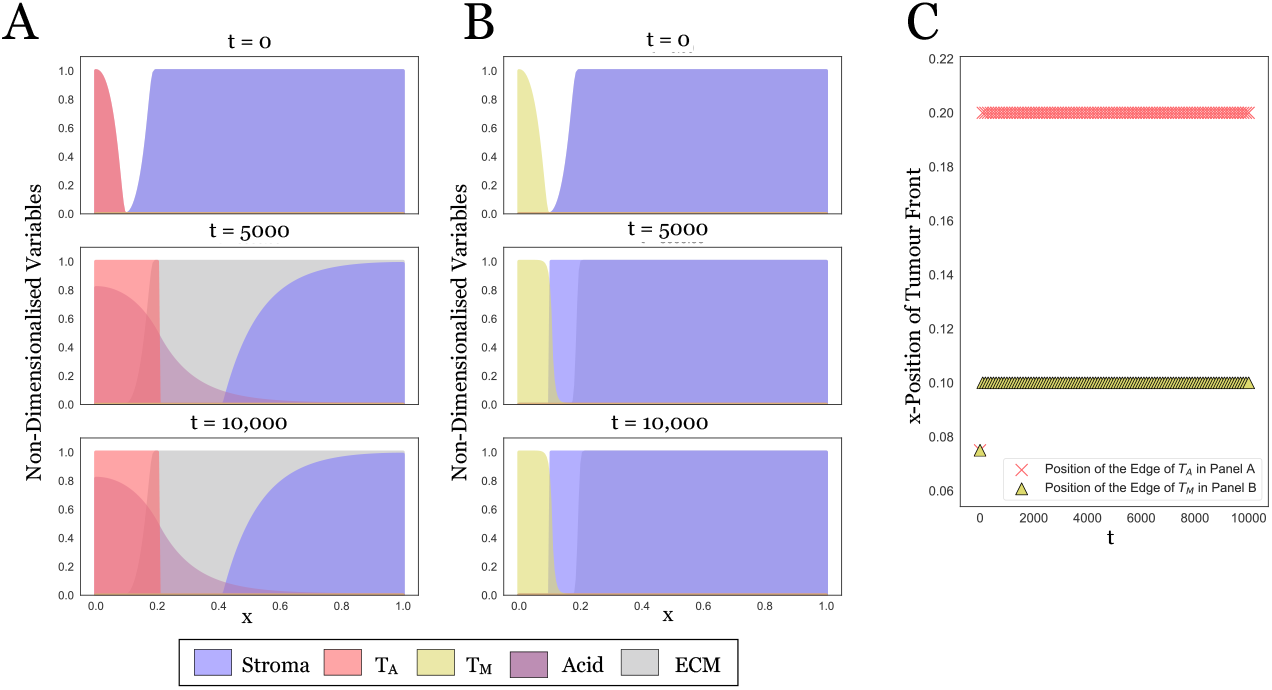
In isolation the tumour populations fail to invade. A: Snapshots at three time points from a long term simulation (*t*_end_ = 10, 000, corresponding to more than 10 years) in which only *T*_*A*_ is present. Expansion stalls because of obstruction by the matrix. B: Analogous simulation of the dynamics with *T*_*M*_ in isolation. This time the tumour can not overcome the stroma. C: Plot showing the position of the tumour edge in A and B over time, determined as min_*x*∈[0,1]_ *d/dt*(*T*_*i*_(*t*)) for *i* = A, M, respectively. We conclude that the model and the numerical scheme behave as expected and that any invasion seen later in this paper is due to the interaction between the two cell types.

## 4 Numerical Simulations

We simulate our model using the method of lines by discretizing space, and then applying a standard ODE integration scheme in time. We discretise the equations in space using the following central difference scheme:

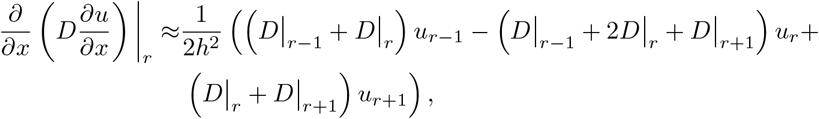

where |_*r*_ denotes evaluation at the r^th^ spatial grid point, *x*_*r*_, of an equi-spaced grid with grid size *h*. In the case of the standard Laplacian operator (as in (11)) this reduces to the standard three-point stencil, whereas for (9)–(10) it provides a consistent discretization of the nonlinear diffusive flux due to the presence of the matrix *M*. The resulting system of ODEs is solved with backwards differentiation formulas (BDF1-BDF5) [48] implemented in Scipy (specifically, the scipy.integrate.ode class). To improve numerical stability, a stabilisation scheme is used to guide state variables back to zero should they become negative. Convergence in space and time for this scheme was checked thoroughly (not shown). The solutions presented are at a resolution of *δx* = 5 × 10^−3^ in space (200 equally spaced points) and relative and absolute numerical tolerances of 1 × 10^−10^ were used for the solution in time. Unless otherwise stated, simulations were run until time *t* = 50 (corresponding to around 575 days). All simulations were carried out in Python 3.6, using Scipy 1.1.0 and Numpy 1.15.1. Visualisations were produced with Pandas 0.23.4, Matplotlib 2.2.3, and Seaborn 0.9.0. The code will be made available upon acceptance of the manuscript.

### 4.1 Neither acid-producing nor matrix-degrading tumour cells invade in isolation

In Figure 3 we show model simulations in which only one of the two populations is present. We see that in isolation neither *T*_*A*_ nor *T*_*M*_ can invade. In accordance with the linear analysis in Section 3, we see that if only *T*_*A*_ is present, then the tumour initially advances but invasion halts because of obstruction by the matrix (Figure 3A). Similarly, if only *T*_*M*_ is present, then the tumour is encapsulated by the stroma (Figure 3B). Plotting the position of the tumour edge in each case confirms this (Figure 3C).

### 4.2 Intra-tumoural competition determines the tumour’s invasion properties

Our results in Section 3 show that when both tumour cell populations are present there are four different possible outcomes depending on the strength of the inter-species competition between *T*_*A*_ and *T*_*M*_. To further investigate this relationship we simulated invasion for 10^4^ combinations of values of (*c*_*M,A*_, *c*_*A,M*_) equally spaced on the grid [0, 2] × [0, 2], corresponding to rates of inter-species competition between 0- and 2-fold that of the intra-species competition. We initialised the tumour as described in Section 2.3 with *σ* = 0.2 and *ω* = 0.1 and simulated until time *t* = 50.

#### 4.2.1 Stable coexistence of multiple tumour phenotypes promotes invasion

Figure 4A shows the position of the tumour edge at *t* = 50 for these 10^4^ parameter combinations. We find that the tumour invades furthest for *c*_*M,A*_, *c*_*A,M*_ < 1, corresponding to the case when inter-species competition is weaker than intra-species competition. Studying the solution for (*c*_*M,A*_, *c*_*A,M*_) = (0, 0) shows that for this range of values the two populations mix and advance as a single front (Figure 5A).

**Fig. 4.**
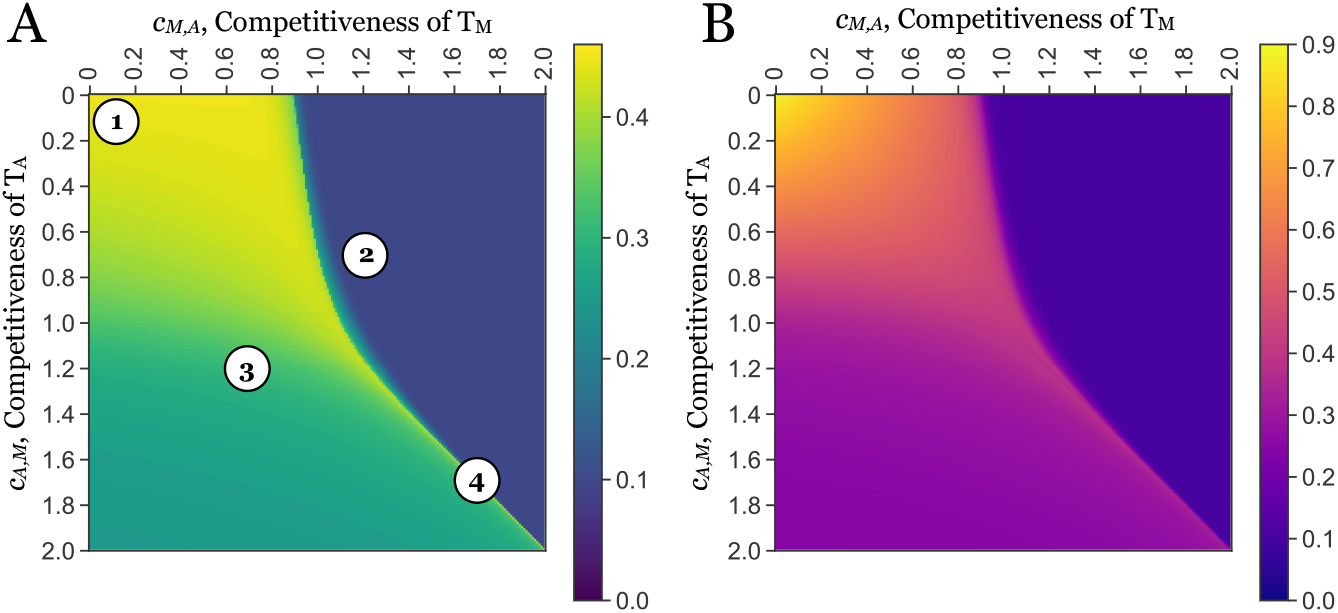
The invasive potential of a tumour is determined by the competition between its subpopulations. A: Position of the tumour front at time t = 50 (575 days) as a function of the strength of inter-species competition (*c*_*M,A*_ and *c*_*A,M*_). This was defined as max(*x*_*A*_, *x*_*M*_), where *x*_*i*_ = min_*x*∈[0,1]_ *{d/dt*(*T*_*i*_(50))} for *i* = *A* or *M*, respectively, is the position of the wave front of *T*_*A*_ and *T*_*M*_ at time 50. Annotations correspond to the time series plots shown in Figure 5. We find the tumour advances furthest when inter-species competition is weaker than intra-species competition (*c*_*M,A*_, *c*_*A,M*_ < 1). B: Total tumour mass, defined as 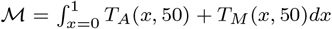, as a function of the inter-species competition. We see that the total tumour mass in the invading tumour, which may be interpreted as a proxy for the total cell number, is a strictly and rapidly decreasing function of the competition parameters. Thus, competition between tumour cells influences not only how far they invade, but also how many cells make up the advancing tumour. Note: cases in which the cell populations were small 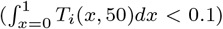 were disregarded in this analysis to avoid issues associated with the simulation and interpretation of low densities.

**Fig. 5.**
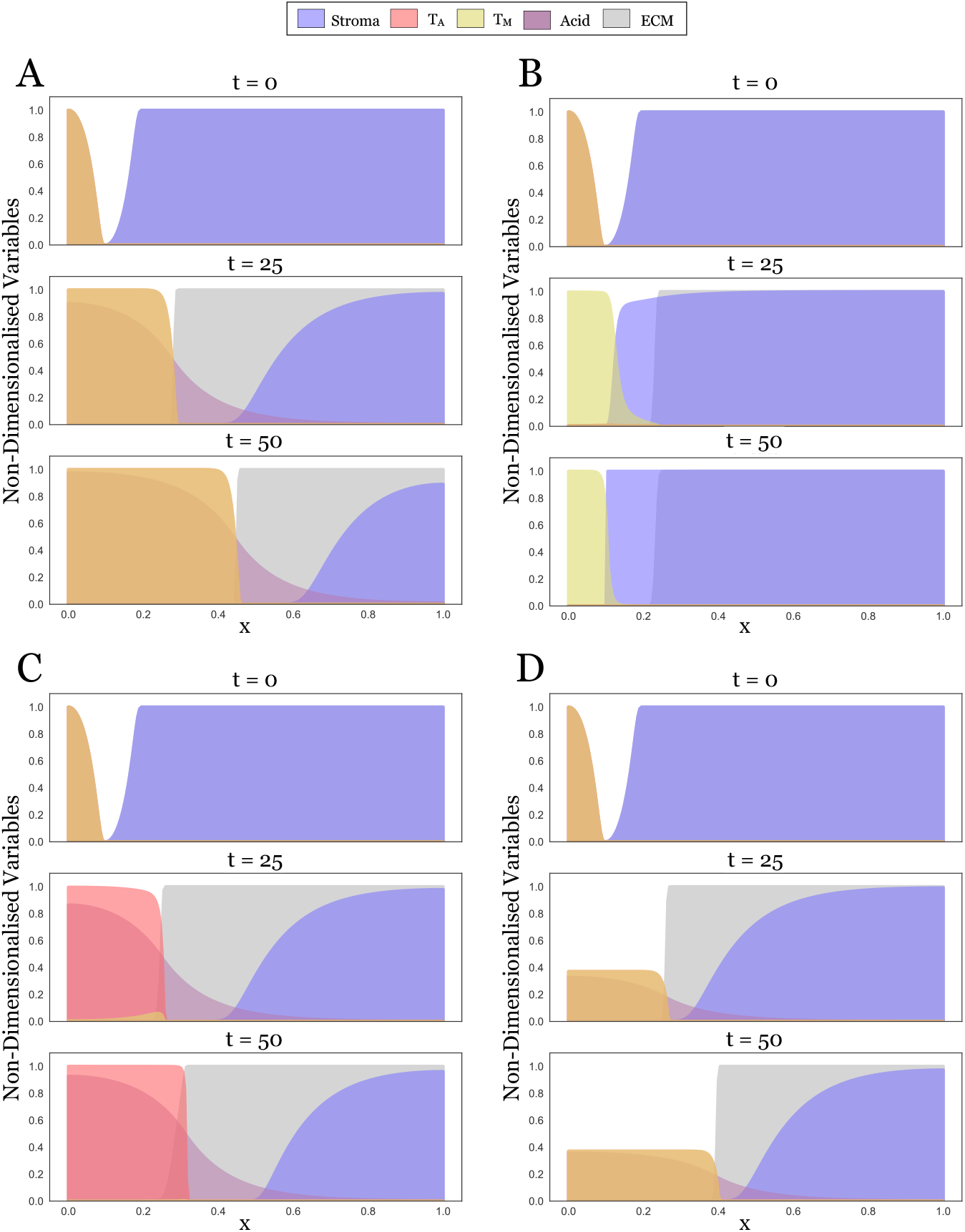
Simulations illustrating the four different scenarios that can occur depending on the inter-species competition between *T*_*A*_ and *T_M_*. Panels correspond to the locations in the competition parameter space marked in Figure 4A (1:A, 2:B, 3:C, 4:D). A: (*c*_*M,A*_, *c*_*A,M*_) = (0, 0). Both tumour populations coexist and together invade rapidly. B: (*c*_*M,A*_, *c*_*A,M*_) = (1.2, 0.7). *T*_*M*_ drives *T*_*A*_ to extinction and invasion halts. C: (*c*_*M,A*_, *c*_*A,M*_) = (0.7, 1.2). *T*_*A*_ dominates over *T*_*M*_. While invasion eventually stops due to a lack of ECM degradation, the tumour initially invades thanks to a small population of *T_M_* persisting at the tumour edge. D: (*c*_*M,A*_, *c*_*A,M*_) = (1.7, 1.7). Mutual exclusion of *T*_*A*_ and *T*_*M*_. When seeded at equal densities the two populations will invade as shown, but the invading front is not stable. If a small perturbation is introduced the two populations will separate and invasion will halt (Figure A2).

Furthermore we observe that the relationship between the invaded distance and the competition parameters is not symmetric about *c*_*M,A*_ and *c*_*A,M*_. In particular, provided *c*_*M,A*_, *c*_*A,M*_ < 1 the invaded distance is more sensitive to a higher competitiveness of *T*_*A*_ than *T*_*M*_. Repeating the experiment in Figure 4A with different rates of matrix degradation, *κ*, shows that this asymmetry is due to *κ* (Appendix A3). For the parameters shown in Figure 4A matrix remodelling is less effective than removal of the stroma for the parameters shown, essentially creating a bottleneck. Our results in Section 3 show that a larger ratio of *c*_*M,A*_ to *c*_*A,M*_ corresponds to a larger proportion of *T*_*M*_ in steady state allowing for more matrix degradation to take place. To summarise, we find that the most invasive tumours are those in which *T*_*A*_ and *T*_*M*_ mix and locally coexist in the correct proportions.

#### 4.2.2 Competitive exclusion slows tumour invasion

As *c*_*M,A*_ is increased through 1, so that *c*_*M,A*_ > 1 and *c*_*A,M*_ < 1, we observe a rapid reduction in tumour expansion (Figure 4A). A simulation for (*c*_*M,A*_, *c*_*A,M*_) = (1.2, 0.7) shows that in this domain, *T*_*M*_ drives *T*_*A*_ to extinction inside the tumour, and is subsequently encapsulated by the stroma due to a lack of acid to keep the stroma in check (Figure 5B).

Similarly, if *T*_*A*_ out-competes *T*_*M*_ (*c*_*M,A*_ < 1 and *c*_*A,M*_ > 1), then invasion is also reduced (Figure 4A). However, this reduction is less significant than in the converse case. This is because the *T*_*M*_ population transiently survives near the edge of the tumour (t = 25 in Figure 5C), where it degrades the ECM for the advancing bulk of the tumour until it is eventually eradicated (t = 50 in Figure 5C and Figure A4).

#### 4.2.3 Strong inter-species competition prevents clonal mixing and reduces invasion

When inter-species competition is stronger than intra-species competition for both populations (*c*_*M,A*_, *c*_*A,M*_ > 1), we observe three possible outcomes: i) the two populations co-exist and invade (*c*_*M,A*_ = *c*_*A,M*_ in Figure 4A), ii) *T*_*A*_ out-competes *T_M_* and the tumour advances only temporarily (*c*_*M,A*_ < *c*_*A,M*_ in Figure 4A), and iii) *T*_*M*_ out-competes *T*_*A*_ and invasion rapidly halts (*c*_*M,A*_ > *c*_*A,M*_ in Figure 4A).

When invasion does occur (*c*_*M,A*_ = *c*_*A,M*_), the tumour is also a mixture of the two phenotypes (Figure 5D), however, the advancing front is unstable to small perturbations (Figure A2). Similarly, if the two populations are not initialised identically, but placed slightly apart, they separate spatially (Figure A2B). Moreover, the solution is strongly sensitive to the parameters, with slight perturbations generating qualitatively different outcomes from the same initial conditions (Figures A2C & D). In summary, this indicates that cooperation in this regime is unstable, and most likely competitive exclusion or spatially separated populations (parapatry) would be observed.

Formally speaking, solutions along the line in the (*c*_*M,A*_, *c*_*M,A*_) parameter space are structurally unstable, corresponding to a separatrix between competitive exclusion of each species. Specifically, away from the invasion front, the stroma, matrix, and acid can be neglected, and the system is simply two Lotka-Volterra-type equations with identical parameters. Neglecting the spatial dynamics and considering the phase-plane of such a system, we see that the stable manifold of the coexistence steady state (which is a saddle) forms a separatrix between the single-species equilibria (see, for instance, Chapter 3 of [39]). This implies that any asymmetry in the initial condition between these two species will lead to one or the other species becoming extinct. Spatial dynamics can then lead to a stabilization of local equilibria of each species, but not to any kind of homogeneous coexistence equilibria, and the spatial structure of the populations can depend sensitively on the initial data. We remark that this separatrix exists even for distinct competition parameters, but for comparable initial densities we do not observe coexistence.

#### 4.2.4 The ratio of invaded distance to tumour mass reflects tumour ecology

In addition to the distance the tumour has invaded, another important feature in the clinic is the total tumour mass that has developed. We compute this as 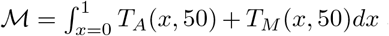 and present the results in Figure 4B. This shows that the two measures are not identical. While the progress of the front is almost identical along the line (*c*_*M,A*_, *c*_*A,M*_) = (*s*, 0) for *s* ∈ [0, 1] (Figure 4A), the mass of the resulting tumour decreases rapidly (Figure 4B). A similar pattern holds true along the line *c*_*M,A*_ = *c*_*A,M*_, and suggests that the strength of competition between tumour sub-populations affects not only the speed of invasion, but also the density of the resulting tumour mass.

## 5 Discussion

While tumour heterogeneity is now widely recognised [38, 29, 3], we are only beginning to comprehend its implications for cancer progression. The fact that the cells in a tumour are not identical, and instead might act as a collective composed of phenotypically distinct individuals, is particularly important in the context of cancer invasion. Invasion of tissue requires both the ability to degrade or remodel the ECM, and the ability to remove surrounding stromal cells. While over time it is possible for the necessary genetic changes to all accumulate in one tumour cell type, it seems more likely that these abilities initially arise in separate cells. Here we aimed to investigate whether cooperation between distinct phenotypic populations is a viable mechanism for invasion, and to characterise the dynamics of such cooperative invasion.

We summarise our findings in Figure 6. Our theoretical results show that cooperation between two cell types gives rise to an invading tumour at clinically realistic speeds (1-2cm in a year). Further, we identify two possible modes of invasion: Firstly, when the two cell types compete weakly with each other, allowing both to stably co-exist. Our model predicts that the resulting mutualistic community has strong invasive potential, as all required traits are present in the same place at the same time (Figure 5A). It has previously been observed that tumours with high degrees of clonal mixing are more aggressive [45, 55]. This has so far been explained by higher cell motility and resulting invasive potential [45]. Based on our results, we propose that an additional explanation could be that mixing allows individual cells to more readily share their traits. As such we advocate further research into the clinical importance of clonal mixing.

**Fig. 6.**
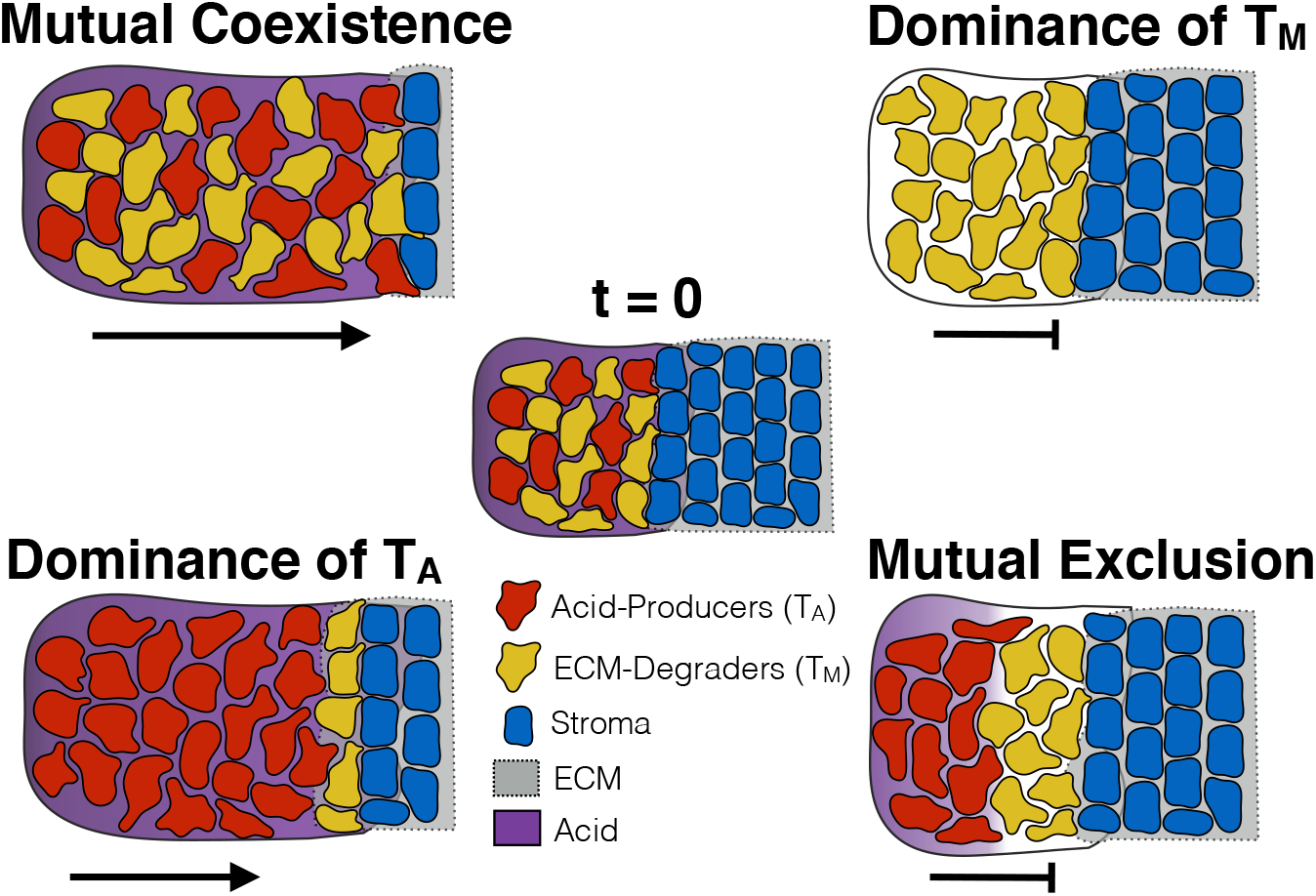
Summary of the key findings of this paper. If the two phenotypes can coexist, a highly invasive community of cells emerges. Conversely, if *T*_*M*_ dominates, tumour invasion comes to a halt as the cells are unable to overcome the stroma. If *T*_*A*_ dominates, a temporarily invasive tumour mass forms in which *T*_*M*_ cells find a temporary habitat in the matrix at the tumour edge. Finally, in the case where the two cell types mutually exclude each other’s growth, the cells separate into spatially distinct regions and fail to invade.

In addition, our model predicts a second mode of invasion, in which the acid producing cells drive the matrix degrading cells to extinction throughout the tumour, but can temporarily invade as a population of matrix degrading cells transiently survives near the edge of the tumour (Figure 6). While invasion in this case is only transient, it could be a contributing factor to cancer invasion, since further mutations could develop or blood vessels could be reached that would allow for continued growth. Current literature suggests that acid producing cells would have a competitive advantage over matrix degrading cells since they are better adapted to low pH conditions [26, 25, 27], and that the onset of invasion is marked by the expansion of a highly glycolytic cancer phenotype [44]. Our results indicate that commensualistic or parasitic relationships might develop between aggressive glycolytic cells in the core of the tumour and cells at the tumour edge which might facilitate invasion. Mathematically, our work also illustrates recent results showing that if the dominant species in a diffusive Lotka-Volterra system moves at a slower rate, then the two species invade empty space as a “propagating-terrace”, where the weaker species invades first but is subsequently eradicated by the dominant species [13].

Although it was not our primary objective, our work also highlights the differences between physical and biological barriers to tumour invasion. In our model, the ECM was a purely physical barrier, whereas the stroma acted by suppressing tumour growth. Figure 3 shows that the biological barrier of the stroma is more effective in blocking tumour invasion than the “wall” of ECM. Unless the level of the ECM is precisely 1, *T*_*A*_ can invade even in the absence of matrix degrading activity and advances until *x* = 0.3 (Figure 3A). In contrast, *T*_*M*_ is stopped at *x* = 0.2 because the arriving tumour cells fail to establish a locally self-sustaining population due to the growth inhibition by the stroma (Figure 3B). This makes the point that a key challenge for invading tumour cells is to survive and grow in this new environment. Furthermore, we found in modelling obstruction that there remains a number of unsolved mathematical challenges: i) How do the travelling wave solutions to this nonlinear diffusion model of movement obstruction develop (Equations (10) & (12))? ii) How do these compare with alternative models of a hard boundary, such as a moving boundary [19, 20]? iii) How should one model distinct, yet simultaneously acting physical obstructions?

We note that there are a number of potentially important interactions not accounted for in the model. Firstly, we do not model matrix regeneration (e.g. [35]). It seems plausible that matrix regeneration might make it significantly more challenging for the matrix degrading cells to invade. As a result, the invasive capabilities of a tumour with a “pocket” of matrix degrading cells might be much smaller than predicted by our model. Conversely, as we discussed in the introduction, some MDEs generate by-products which can stimulate movement of the cells. Anderson et al [2] found that this can result in the leading edge of the tumour separating from the main mass. In our model this might allow the matrix degrading cells to penetrate further into the tissue and increase invasiveness. Finally, the ECM is composed of proteins and as such also subject to acid degradation [37]. Because the aim of this paper was to acquire a first understanding of what general behaviours might emerge we neglected this degradation in our model. However, clearly, this will influence the invasive behaviour and it would be important to include such a term in future models. Finally, we remark that our approach focused on understanding the invasive front itself, using a simplified model of phenotype interaction (direct competition). It is now well-known that the selection pressures at the edge of an invasive front are different from within an organism’s “home range” due to a range of differences near an invading front (the Allee and Olympic Village effects, for instance) [33, 41, 21, 12]. More generally, evolution and life-history can have strong impacts on dispersal efficiency and range expansion [8, 10, 40]. Investigating these different modes of selection could provide insight into phenotypic heterogeneity throughout a tumour compared to its invading edge.

To sum up, we have explored cooperation of tumour cells as a mode of tumour invasion. We found that the most invasive tumour emerges when cells coexist in the same region in space as this allows cells to most effectively share their traits. This point is simple but important: To fully understand the implications of tumour heterogeneity we have to ask not only *what cells are present* but also *where are these cells located* ? Do they live in separate regions or can they spatially *mix* and, thus, *match* their traits?

## Acknowledgements

We would like to thank Dr Jonathan Whitely and Dr Faustino Sánchez-Garduño for helpful discussions about the simulation and properties of degenerate travelling waves. Further we would like to thank Dr Chandler Gatenbee for his help in processing the images for Figure 1. This research was supported by funding from the Engineering and Physical Sciences Research Council (EPSRC) and the Medical Research Council (MRC) [grant number EP/L016044/1]. Anderson gratefully acknowledges funding from both the Cancer Systems Biology Consortium and the Physical Sciences Oncology Network at the National Cancer Institute, through grants U01CA232382 and U54CA193489 as well as support from the Moffitt Center of Excellence for Evolutionary Therapy.

## A1 Image Collection and Processing

A TMA containing formalin-fixed and paraffin-embedded human breast tissue specimens was constructed at the Moffitt Cancer Center histology core. The TMA contains 27 normal breast tissue, 30 DCIS, 48 invasive ductal carcinomas without metastasis, 49 invasive ductal carcinomas with metastasis and 48 lymph node macrometastases of breast cancer. Cores were selected from viable tumour regions and did not contain necrosis. A 1:200 dilution of anti-LAMP2b (#ab18529, Abcam) and 1:200 of anti-TGM2 (#ab109200, Abcam) were used as primary antibodiy. Normal placenta was used as a positive control for LAMP2 and normal human kidney for TGM2. For the negative control, an adjacent section of the same tissue was stained without application of primary antibody, and any stain pattern observed was considered as nonspecific binding of the secondary.

Immunohistochemical analysis was conducted using digitally scanning slides. A pathologist reviewer scored the intensity of each stain on a scale from 0 to 3, where a 0 was considered negative, score 1 was weakly positive, score 2 was moderately positive and score 3 was strongly positive. For further information see [17].

In order to create the visualisations in Figure 1 we extracted masks of only the areas with the highest score (a score of 3). To do so we first aligned the slides for each stain (LAMP2b and TGM2) using VALIS (in preparation; see also [23]), and subsequently extracted the areas with the highest score using OpenCV. Finally we overlaid the extracted masks on top of the TGM2 slide.

## A2 Stability Analysis

In Table A1, we list the eigenvalues computed for each steady state. For SS4 and SS6 no simple analytic forms were obtainable. Instead, we numerically computed the eigenvalues for the range of parameters of interest and show 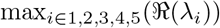, where *λ*_*i*_ denotes the i^th^ eigenvalue of the Jacobian (Figure A1). We conclude that only SS0, SS1, and SS3 are stable. Analytic results were obtained by hand and confirmed with Maple 2018. Numerical computations were carried out in Python 3.6 (for further details on the environment see Section 4).

## A3 Sensitivity Analysis for *κ* and *c*_*S*_

As the parameters for the strength of competition between the stroma and the tumour (*c*_*S*_) and the rate of matrix degradation are difficult to obtain experimentally, we performed a sensitivity analysis over the plausible range in which they might lie. In Figure A3 we show the distance invaded by the tumour, calculated as for Figure 4. We see that as *κ* is increased the tumour invades further. Moreover, whilst for values of *κ* = 10 tumours with higher proportion of *T*_*M*_ (corresponding to lower *c*_*A,M*_; see also Figure 4A) appear more aggressive, this is less apparent when the matrix can be degraded faster (*κ* = 100).

---

**Table A1.**
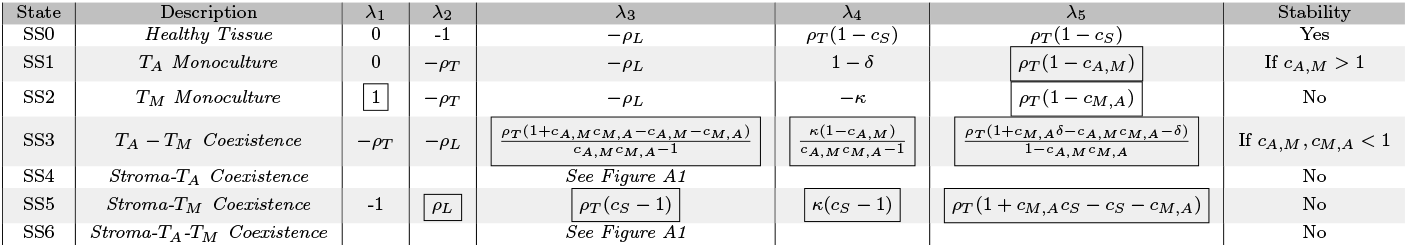
The eigenvalues describing the linear stability of the steady states. Stability is assessed based on the parameter values in Table 1. Boxed values indicate eigenvalues that are positive in at least parts of the parameters space. For SS4 and SS6 no simple analytic forms were attainable, and instead stability was assessed numerically (see Figure A1).

**Fig. A1.**
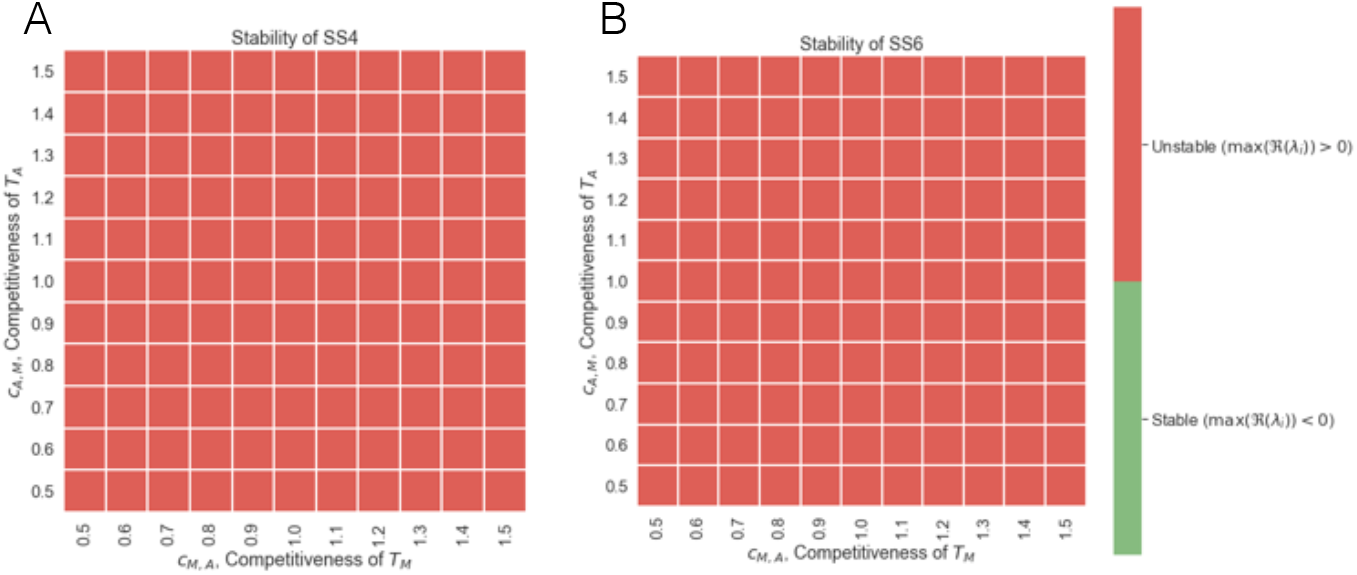
Numerical stability analysis of SS4 (A) and SS6 (B) for the range of parameters considered in this paper (see Table 1). Stability was assessed by computing the eigenvalues of the Jacobian at the steady state, and assessing whether at least one eigenvalue had a strictly positive real part. We find that both SS4 and SS6 are unstable across the range of parameters considered.

**Fig. A2.**
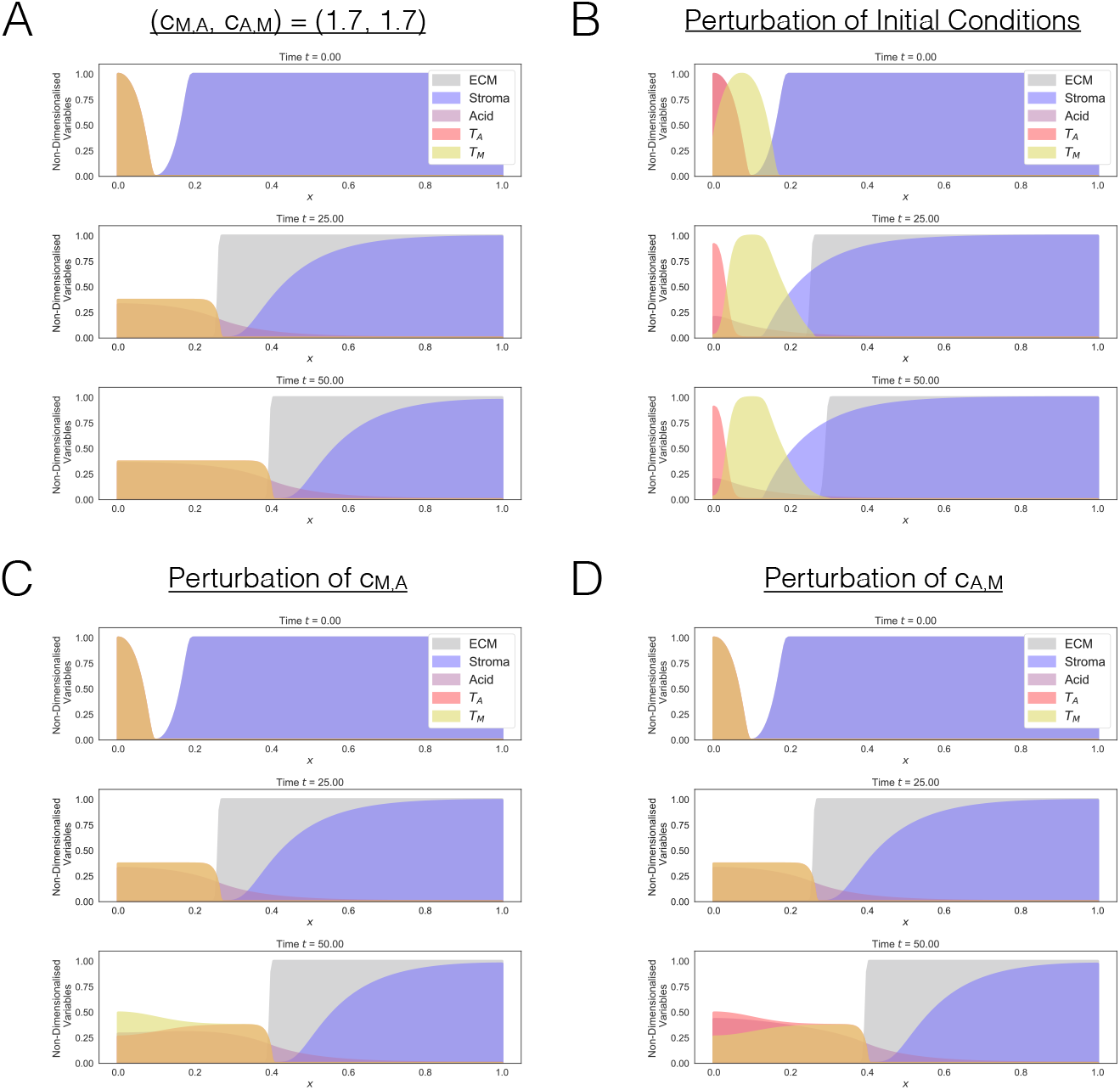
Instability of the invasive front when *c*_*M,A*_ = *c*_*A,M*_. A: Simulation for *c*_*M,A*_ = *c*_*A,M*_ = 1.7. B: Perturbation of the initial conditions in A. The initial distribution of *T*_*M*_ was shifted by *δx* = 0.15 to the right, resulting in subsequent spatial separation of the two phenotypes and arresting of invasion. C: Small perturbation in *c*_*M,A*_ ((*c*_*M,A*_, *c*_*A,M*_) = (1.700001, 1.7)). D: Small perturbation in *c*_*A,M*_ ((*c*_*M,A*_, *c*_*A,M*_) = (1.7, 1.700001)). This indicates that the system is structurally unstable in this parameter regime.

**Fig. A3.**
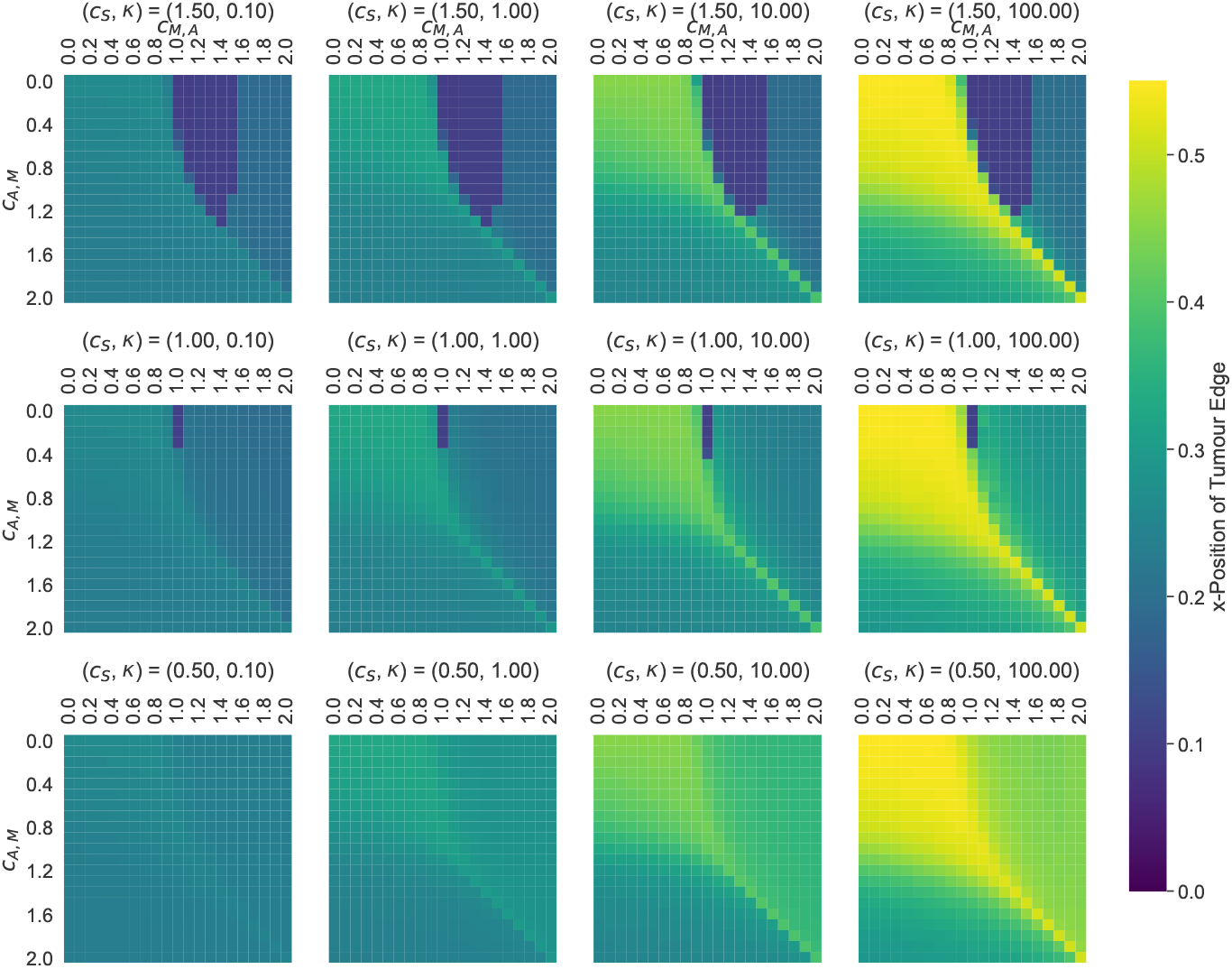
Sensitivity analysis for the parameters *c*_*M,A*_, *c*_*A,M*_, *c*_*S*_ and *κ*. Each heatmap shows the position of the front of the tumour at t = 50 (computed as for Figure 4A).

**Fig. A4.**
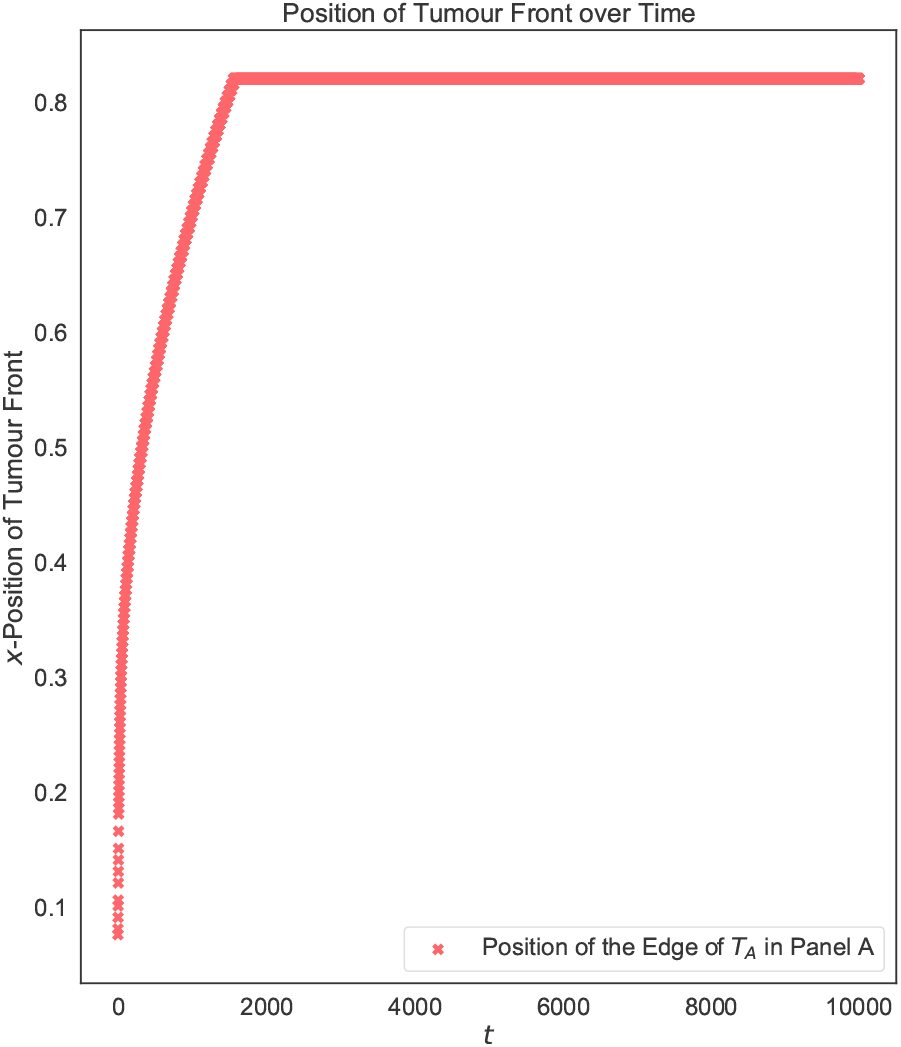
When *T*_*A*_ drives *T*_*M*_ to extinction only transient invasion occurs. Position of the tumour edge simulated until *t* = 10000 for (*c*_*M,A*_, *c*_*A,M*_) = (0.5, 1.5). The tumour transiently advances while the *T*_*M*_ initially persists. However, eventually *T*_*M*_ is eradicated and invasion stalls.

## References

1. Andasari, V., Gerisch, A., Lolas, G., South, A.P., Chaplain, M.A.J.: Mathematical modeling of cancer cell invasion of tissue: Biological insight from mathematical analysis and computational simulation. Journal of Mathematical Biology 63(1), 141–171 (2011). DOI 10.1007/s00285-010-0369-1. URL https://link.springer.com/content/pdf/10.1007%2Fs00285-010-0369-1.pdf

2. Anderson, A.R.A., Chaplain, M.A.J., Newman, L.E., Steele, R.J.C., Thompson, A.M.: Mathematical modelling of tumour invasion and metastasis. Computational and mathmatical methods in medicine 2(2), 129–154 (2000)

3. Anderson, A.R.A., Maini, P.K.: Mathematical Oncology. Bullet of Mathematical Biology 80(5), 945–953 (2018). DOI 10.1007/s11538-018-0423-5. URL http://link.springer.com/10.1007/s11538-018-0423-5

4. Araujo, R.P., McElwain, D.L.S.: A history of the study of solid tumour growth: The contribution of mathematical modelling. Bulletin of Mathematical Biology 66(5), 1039–1091 (2004). DOI 10.1016/j.bulm.2003.11.002. URL https://www.sciencedirect.com/science/article/pii/S0092824003001265

5. Archetti, M., Pienta, K.J.: Cooperation among cancer cells: applying game theory to cancer. Nature Reviews Cancer p. 1 (2018). DOI 10.1038/s41568-018-0083-7. URL http://www.nature.com/articles/s41568-018-0083-7

6. Axelrod, R., Axelrod, D.E., Pienta, K.J.: Evolution of cooperation among tumor cells. Proceedings of the National Academy of Sciences 103(36), 13474–13479 (2006)

7. Basanta, D., Anderson, A.R.A.: Exploiting ecological principles to better understand cancer progression and treatment. Interface Focus 3(4), 20130020 (2013)

8. Benichou, O., Calvez, V., Meunier, N., Voituriez, R.: Front acceleration by dynamic selection in fisher population waves. Physical Review E 86(4), 041908 (2012)

9. Bloom, A.B., Zaman, M.H.: Influence of the microenvironment on cell fate determination and migration. Physiological genomics 46(9), 309–314 (2014)

10. Bouin, E., Calvez, V., Meunier, N., Mirrahimi, S., Perthame, B., Raoul, G., Voituriez, R.: Invasion fronts with variable motility: phenotype selection, spatial sorting and wave acceleration. Comptes Rendus Mathematique 350(15-16), 761–766 (2012)

11. Byrne, H.M., Chaplain, M.A.: Modelling the role of cell-cell adhesion in the growth and development of carcinomas. Mathematical and Computer Modelling 24(12), 1–17 (1996). DOI 10.1016/S0895-7177(96)00174-4. URL https://www.sciencedirect.com/science/article/pii/S0895717796001744

12. Calvez, V., Henderson, C., Mirrahimi, S., Turanova, O., Dumont, T.: Non-local competition slows down front acceleration during dispersal evolution. arXiv preprint arXiv:1810.07634 (2018)

13. Carrère, C.: Spreading speeds for a two-species competition-diffusion system. Journal of Differential Equations 264(3), 2133–2156 (2018). DOI 10.1016/J.JDE.2017.10.017. URL https://www.sciencedirect.com/science/article/pii/S0022039617305612

14. Chaplain, M.A.J., Lolas, G.: Mathematical modelling of cancer cell invasion of tissue: The role of the urokinase plasminogen activation system. Mathematical Models and Methods in Applied Sciences 15(11), 1685–1734 (2005). DOI 10.1142/S0218202505000947. URL https://www.worldscientific.com/doi/abs/10.1142/S0218202505000947

15. Cleary, A.S., Leonard, T.L., Gestl, S.A., Gunther, E.J.: Tumour cell heterogeneity maintained by cooperating subclones in Wnt-driven mammary cancers. Nature 508(7494), 113–117 (2014). DOI 10.1038/nature13187. URL http://www.nature.com/articles/nature13187

16. Curran, S., Murray, G.I.: Matrix metalloproteinases in tumour invasion and metastasis. The Journal of pathology 189(3), 300–308 (1999)

17. Damaghi, M., Byrne, S., Xu, L., Tafreshi, N., Fang, B., Koomen, J.M., Karolak, A., Chen, T., Johnson, J., Gallant, N.D., Marusyk, A., Gillies, R.J.: Collagen Production and Niche Engineering: A Novel Strategy for Cancer Cells to Survive Acidosis and Evolve. bioRxiv p. 711978 (2019). DOI 10.1101/711978. URL https://www.biorxiv.org/content/10.1101/711978v1.abstract

18. Domschke, P., Trucu, D., Gerisch, A., A. J. Chaplain, M.: Mathematical modelling of cancer invasion: Implications of cell adhesion variability for tumour infiltrative growth patterns. Journal of Theoretical Biology 361, 41–60 (2014). DOI 10.1016/J.JTBI.2014.07.010. URL https://www.sciencedirect.com/science/article/pii/S0022519314004032

19. Du, Y., Guo, Z.: The stefan problem for the fisher–kpp equation. Journal of Differential Equations 253(3), 996–1035 (2012)

20. El-Hachem, M., McCue, S.W., Jin, W., Du, Y., Simpson, M.J.: Revisiting the fisher-kpp equation to interpret the spreading-extinction dichotomy. boiRxiv p. 673202 (2019)

21. Erm, P., Phillips, B.L.: Evolution transforms pushed waves into pulled waves. bioRxiv p. 266007 (2018)

22. Fasano, A., Herrero, M.A., Rodrigo, M.R.: Slow and fast invasion waves in a model of acid-mediated tumour growth. Mathematical Biosciences 220(1), 45–56 (2009). DOI 10.1016/j.mbs.2009.04.001. URL http://www.ncbi.nlm.nih.gov/pubmed/19376138

23. Gatenbee, C.D., Baker, A.M., Schenck, R.O., Neves, M.P., Hasan, S.Y., Martinez, P., Cross, W.C., Jansen, M., Rodriguez-Justo, M., Sottoriva, A., Leedham, S., Robertson-Tessi, M., Graham, T.A., Anderson, A.R.: Niche engineering drives early passage through an immune bottleneck in progression to colorectal cancer. bioRxiv (2019). DOI 10.1101/623959. URL https://www.biorxiv.org/content/early/2019/05/03/623959.1

24. Gatenby, R.A., Gawlinski, E.T.: A Reaction-Diffusion Model of Cancer Invasion. Cancer Research 56(31), 5745–5753 (1996). URL http://cancerres.aacrjournals.org/content/56/24/5745.abstract

25. Gatenby, R.A., Gawlinski, E.T., Gmitro, A.F., Kaylor, B., Gillies, R.J.: Acid-mediated tumor invasion: a multidisciplinary study. Cancer research 66(10), 5216–5223 (2006)

26. Gatenby, R.A., Gillies, R.J.: Why do cancers have high aerobic glycolysis? Nature Reviews Cancer 4(11), 891–899 (2004)

27. Gatenby, R.A., Smallbone, K., Maini, P.K., Rose, F., Averill, J., Nagle, R.B., Worrall, L., Gillies, R.J.: Cellular adaptations to hypoxia and acidosis during somatic evolution of breast cancer. British journal of cancer 97(5), 646–653 (2007)

28. Gerisch, A., Chaplain, M.A.J.: Mathematical modelling of cancer cell invasion of tissue: Local and non-local models and the effect of adhesion. Journal of Theoretical Biology 250(4), 684–704 (2008). DOI 10.1016/j.jtbi.2007.10.026

29. Gerlinger, M., Rowan, A.J., Horswell, S., Larkin, J., Endesfelder, D., Gronroos, E., Martinez, P., Matthews, N., Stewart, A., Tarpey, P., Others: Intratumor heterogeneity and branched evolution revealed by multiregion sequencing. New England journal of medicine 366(10), 883–892 (2012)

30. Gillies, R.J., Robey, I., Gatenby, R.A.: Causes and consequences of increased glucose metabolism of cancers. Journal of Nuclear Medicine 49(2), 24S (2008)

31. Hanahan, D., Weinberg, R.A.: The hallmarks of cancer. cell 100(1), 57–70 (2000)

32. Helmlinger, G., Yuan, F., Dellian, M., Jain, R.K.: Interstitial pH and pO2 gradients in solid tumors in vivo: high-resolution measurements reveal a lack of correlation. Nature Medicine 3(2), 177 (1997)

33. Keymer, J.E., Marquet, P.A.: The complexity of cancer ecosystems. Frontiers in Ecology, Evolution and Complexity pp. 101–119 (2014)

34. Kim, S., Goel, S., Alexander, C.M.: Differentiation Generates Paracrine Cell Pairs That Maintain Basaloid Mouse Mammary Tumors: Proof of Concept. PLoS ONE 6(4), e19310 (2011). DOI 10.1371/journal.pone.0019310. URL http://dx.plos.org/10.1371/journal.pone.0019310

35. Martin, N.K., Gaffney, E.A., Gatenby, R.A., Maini, P.K.: Tumour–stromal interactions in acid-mediated invasion: a mathematical model. Journal of Theoretical Biology 267(3), 461–470 (2010)

36. McGillen, J.B., Gaffney, E.A., Martin, N.K., Maini, P.K.: A general reaction–diffusion model of acidity in cancer invasion. Journal of Mathematical Biology 68(5), 1199–1224 (2014)

37. McKinnell, R.G.: The biological basis of cancer. Cambridge University Press (1998)

38. Merlo, L.M.F., Pepper, J.W., Reid, B.J., Maley, C.C.: Cancer as an evolutionary and ecological process. Nature Reviews Cancer 6(12), 924–935 (2006)

39. Murray, J.D.: Mathematical Biology I. An introduction, 3rd edn. Springer-Verlag, Berlin Heidelberg (2002). DOI 10.1007/b98868

40. Perkins, A.T., Phillips, B.L., Baskett, M.L., Hastings, A.: Evolution of dispersal and life history interact to drive accelerating spread of an invasive species. Ecology Letters 16(8), 1079–1087 (2013)

41. Perkins, T.A., Boettiger, C., Phillips, B.L.: After the games are over: life-history trade-offs drive dispersal attenuation following range expansion. Ecology and evolution 6(18), 6425–6434 (2016)

42. Perumpanani, A., Byrne, H.: Extracellular matrix concentration exerts selection pressure on invasive cells. European Journal of Cancer 35(8), 1274–1280 (1999)

43. Ramis-Conde, I., Chaplain, M.A., Anderson, A.R.: Mathematical modelling of cancer cell invasion of tissue. Mathematical and Computer Modelling 47(5-6), 533–545 (2008). DOI 10.1016/j.mcm.2007.02.034. URL https://linkinghub.elsevier.com/retrieve/pii/S0895717707001823

44. Robertson-Tessi, M., Gillies, R.J., Gatenby, R.A., Anderson, A.R.A.: Impact of metabolic heterogeneity on tumor growth, invasion, and treatment outcomes. Cancer Research 75(8), 1567–1579 (2015)

45. Sottoriva, A., Kang, H., Ma, Z., Graham, T.A., Salomon, M.P., Zhao, J., Marjoram, P., Siegmund, K., Press, M.F., Shibata, D., Curtis, C.: A Big Bang model of human colorectal tumor growth. Nature Genetics 47(3), 209–216 (2015). DOI 10.1038/ng.3214. URL http://www.nature.com/doifinder/10.1038/ng.3214

46. Sporn, M.B.: The war on cancer. The Lancet 347(9012), 1377–81 (1996). URL http://www.ncbi.nlm.nih.gov/pubmed/8637346

47. Stetler-Stevenson, W.G., Aznavoorian, S., Liotta, L.A.: Tumor cell interactions with the extracellular matrix during invasion and metastasis. Annual Review of Cell Biology 9(1), 541–573 (1993)

48. Süli, E., Mayers, D.F.: An introduction to numerical analysis. Cambridge university press (2003)

49. Tannock, I.F., Rotin, D.: Acid pH in tumors and its potential for therapeutic exploitation. Cancer Research 49(16), 4373–4384 (1989)

50. Wadlow, R.C., Wittner, B.S., Finley, S.A., Bergquist, H., Upadhyay, R.: Systems-Level Modeling of Cancer-Fibroblast Interaction. PLoS ONE 4(9), 6888 (2009). DOI 10.1371/journal.pone.0006888

51. Warburg, O.H., Dickens, F.: The Metabolism of Tumors (English translation by F. Dickens). Constable, London (1930)

52. Webb, S.D., Sherratt, J.A., Fish, R.G.: Alterations in proteolytic activity at low ph and its association with invasion: a theoretical model. Clinical & Experimental Metastasis 17(5), 397–407 (1999)

53. Werb, Z.: ECM and cell surface proteolysis: regulating cellular ecology. Cell 91(4), 439–442 (1997)

54. Wike-Hooley, J., Haveman, J., Reinhold, H.: The relevance of tumour ph to the treatment of malignant disease. Radiotherapy and Oncology 2(4), 343–366 (1984)

55. Zhang, A.W., Mcpherson, A., Milne, K., Holt, R.A., Nelson, B.H., Shah, S.P., Kroeger, D.R., Hamilton, P.T., Miranda, A., Funnell, T., Little, N., De Souza, C.P.E., Laan, S., Ledoux, S., Cochrane, D.R., Lim, J.L.P., Yang, W., Roth, A., Smith, M.A., Ho, J., Tse, K., Zeng, T., Shlafman, I., Mayo, M.R., Moore, R., Failmezger, H., Heindl, A., Wang, Y.K., Bashashati, A., Grewal, D.S., Brown, S.D., Lai, D., Wan, A.N.C., Nielsen, C.B., Huebner, C., Tessier-Cloutier, B.: Interfaces of Malignant and Immunologic Clonal Dynamics in Ovarian Cancer. Cell 173(1), 1–15 (2018). DOI 10.1016/j.cell.2018.03.073. URL https://doi.org/10.1016/j.cell.2018.03.073

